# Age and neuroinflammation are important components of the mechanism of cognitive and neurobehavioral deficits in sickle cell disease

**DOI:** 10.1101/2020.03.24.006221

**Authors:** Raven A. Hardy, Noor Abi Rached, Jayre A. Jones, David R. Archer, Hyacinth I. Hyacinth

## Abstract

**Objectives:** Cognitive and neurobehavioral abnormalities are the most common and complex complications of sickle cell disease (SCD). Known risk factors influencing abnormalities are stroke and silent cerebral infarcts, but a majority of cases do not have overt cerebral injury and the underlying mechanism is not well understood. This study aims to determine whether sickle cell mice could recapitulate features of cognitive and neurobehavioral impairment observed in sickle cell patients as well as to determine the underlying cellular mechanism of these SCD complications.

**Methods:** Using a longitudinal cross-sectional study design, we evaluated cognition and neurobehavioral deficits as an outcome. Six as well as 13 months old male Townes humanized sickle cell (SS) and matched control (AA) mice were tested. The combination of novel object recognition and fear conditioning tests was employed to measure anxiety/depression, learning and memory. Immunohistochemistry was performed to quantify bone marrow-derived microglia (CD45^+^) and activated microglia (Iba1^+^) in the dentate and peri-dentate gyrus to determine if these factors were potential pathogenic mechanisms associated with cognitive and neurobehavioral abnormalities. We evaluated neurogenesis by measuring 5’Bromodeoxyuridine (BrdU) and doublecortin (DCX) and phenotyped proliferating cells via quantification of glial fibrillary acid protein (GFAP^+^), neuronal nuclei (NeuN^+^), CD45^+^ and Iba1^+^. In addition, Golgi-Cox staining was used to assess neuroplasticity via measurement of dendritic spine density and morphology, as well as dendrite arbors.

**Results:** Compared to matched AA, 13 months old SS mice showed significant evidence of anxiety/depression by the shorter distance traveled as well as thigmotaxis. Additionally, SS mice were significantly less likely to recognize the novel object as well as have a reduced preference for the novel object. There were no significant differences between 6 months old SS and AA. But the difference reappeared after the same mice were aged to 13 months. Aged mice exhibited more anxiety/depression behaviors and thigmotaxis and were less likely to recognize or show a lower percent preference for the novel object compared to aged control (AA) mice. Immunohistochemistry analysis shows that sickle cell (SS) mice had significantly more CD45^+^ and Iba1^+^ activated microglia cells in the dentate and peri-dentate gyrus area compared to AA mice. SS mice also had a significantly lower dendritic spine density compared with controls. Treatment of aged SS mice with minocycline resulted in significant improvement of cognitive and neurobehavioral function compared to matched vehicle-treated SS mice. Also immunohistochemical and histological analysis showed that treated SS mice had significantly fewer CD45^+^ cells and activated microglia in the dentate and peri-dentate gyrus area. Furthermore, there was significant improvement in dendritic spine and dendrite arbor density as well as spine maturation in treated sickle cell mice compared with vehicle-treated sickle cell mice.

**Conclusion:** Taken together these results indicate that age, neuro-inflammation and neuroplasticity, specifically, spine maturation and density, are possible mechanisms underlying cognition deficits in sickle cell disease. These could also be targeted as a potential approach for prevention and or treatment of cognitive and neurobehavioral deficits in SCD.

## Introduction

Cerebrovascular disease is one of the most common and dramatic complications of sickle cell disease (SCD) affecting both children and adults,^1^ with cognitive and neurobehavioral abnormalities as possible sequelae. The prevalence of cognitive and neurobehavioral complications in individuals with SCD have been widely reported.^2-8^ Some studies have suggested that these abnormalities are a result of stroke and/or repeated silent cerebral infarctions (SCIs) observed in children, adolescents, and adults with SCD. ^9, 10^ Other studies have also associated the presence of cognitive and neurobehavioral dysfunction with older age, anemia, overt stroke and silent cerebral infarcts (SCIs), systemic inflammation as well as environmental stressors (low socioeconomic status) in individuals with SCD.^11-14^ However, recent studies indicate that cognitive and neurobehavioral abnormalities were present in individuals with SCD, even in the absence of magnetic resonance imaging (MRI) detectable infarcts or cerebral injury.^2, 8, 14^ Andreotti et al., recently reported that children with SCD, in the absence of MRI evidence of cerebral infarct or injury, scored below the normative mean on cognitive testing. Further, in the same children, there was a significant negative correlation between plasma inflammatory cytokine levels and tests of executive function.^2^ A similar relationship existed in children with asthma, who have high circulating levels of inflammatory cytokines ^2^ suggesting a possible mechanistic role for inflammation and potentially neuroinflammation. Notwithstanding the abundance of epidemiological evidence, the mechanism of onset of cognitive and neurobehavioral dysfunction in SCD, and the underlying cellular and pathobiological mechanisms are largely unknown.^15-18^

The presence of cognitive and neurobehavioral dysfunction in individuals with SCD in the absence of overt cerebral injury ^2, 14^ suggests the involvement of cellular mechanisms that are not detectable on clinical imaging. The mechanisms of cognitive and neurobehavioral dysfunction in SCD in the absence of overt cerebral injury is largely unknown. But in non-sickle cell animals models, studies have shown that changes in the anatomy of neurons and neuroinflammation (localized or as a result of cross-talk between systemic inflammation and central nervous system inflammation), are associated with cognitive and neurobehavioral dysfunction and deficits.^19-24^ Furthermore, several studies have shown that cerebral ischemia results in abnormal remodeling of neuronal dendrites (decreased arbors) and dendritic spines (decrease in number and maturation) and that decreased number of dendritic arbors and spines and proportion of mature dendritic spines correlate with cognitive and neurobehavioral dysfunction.^25-28^ Also, there is evidence to suggest that neuroinflammation is the underlying mechanism of stress-induced neurocognitive dysfunction.^19, 20, 22^ Taken together, the available data suggests that similar mechanisms as outlined above, might be responsible for SCD-related cognitive and neurobehavioral dysfunction. While it has been suggested that the cognitive and neurobehavioral complications observed among individuals with SCD could be attributed to pain and the chronicity of the disease, available studies have shown that individuals with SCD were 20% more likely than those with hemochromatosis and other chronic hematological diseases, to have anxiety, depression or poor educational attainment.^6^ Additionally, adult patients with SCD were more likely to be depressed and anxious after adjusting for pain frequency, pain and disease severity, and frequency of hospitalization.^29, 30^ This body of evidence suggests that SCD is associated with significant cognitive and neurobehavioral impairment that are independent of the individual burden of disease and/or overt central nervous system injury. Our hypothesis is that age-related neuronal remodeling (decreased density of dendrite arbors, dendritic spines and, proportion of mature dendritic spines) and neuroinflammation are potential mechanisms underlying the development of cognitive and neurobehavioral dysfunction in SCD.

## Methods

### Animal preparation

The Institutional Animal Care and Use Committee (IACUC) of Emory University approved this study. All research activities reported here were conducted in accordance with the National Research Council and National Institutes of Health *Guide for the Care and Use of Laboratory Animals 8*^*th*^ *Edition.*^31^ Reporting of methods, results and all aspects of this paper are in accordance with appropriate reporting standards (ARRIVE). Reporting of methods was based on the recommendations from the *Quality of methods reporting in animal models of colitis.*^32^

We utilized a combination of cross-sectional and cross-sectional prospective study designs. We also used the Townes mouse model, a humanized sickle cell (with HbSS) and corresponding control (with HbAA) mouse model (B6;129-Hba^tm1(HBA)Tow^ Hbb^tm2(HBG1,HBB*)Tow^/Hbb^tm3(HBG1,HBB)Tow^/J) throughout this study. Based on a recent report,^33^ we estimated that an average sample size of 15 mice per group wouldl be adequate for the cognitive and neurobehavioral studies. Furthermore, a sample size of 4 mice would be adequate to show statistically significant differences for the histological studies based on our prior published work in sickle cell mice.^34-36^

Mice for this study were breed at the Emory University managed breeding services, weaning at 3 – 4 weeks of age and were housed with a 12-hour light/dark cycle with water and feed provided *ad libitum*. Mice were allowed to age naturally until ready for use in our experiments described below. Although the required sample size for the cognitive and neurobehavioral studies was 15 mice per genotype or treatment group, we started with 20 mice in each group to allow for a 10 – 20% attrition from the mortality, which is usually higher in sickle cell mice compared to controls.

### Study design and overall methods

For this study, we used only male sickle cell mice. We first set out to determine whether aged (13 months old) sickle cell mice compared to age-matched controls show evidence of learning, cognitive and neurobehavioral deficit. For this experiment aged sickle cell mice and matched controls (cohort I) were tested for the presence of learning, cognitive and neurobehavioral deficits when compared to controls. We used the Novel object recognition (NOR) test which consit of habituation and novel object recognition. We later tested another cohort of 15 sickle cell mice and 12 matched controls (cohort II) who were 6 months old at the time of testing. This was to enable us determine whether aging was an important factor in our observation in the first instance. Next, the cohort II mice were allowed to age until they were 13 months old and then retested using the same neurobehavioral paradigm as cohort I. We added an additional learning and memory test paradigm (fear conditioning) that is based on classical Pavlovian conditioning with a goal of ensuring that observations made in NOR experiments were not spurious and replicable by a different learning and memory test paradigm. We also wanted to determine the possible location of the lesion (amygdala vs. hippocampus or both). Based on emerging evidence,^19, 22, 37^ we tested the hypothesis that neuroinflammation might be a mechanism for the observed differences in learning, cognitive and neurobehavioral function between sickle cell mice and controls by assessing cellular markers of neuroinflammation and by blocking neuroinflammation using minocycline. Minocycline, is a highly lipophilic drug capable of crossing the blood-brain barrier,^38^ and was administered to a subset of previously aged mice (cohort II after habituation and NOR) and later to a larger sample of animals (cohort III, N = 15 per genotype or treatment groups). Minocycline has been shown to have a neuroprotective effect in several neurological disorders^39, 40^ and its mechanism of action is proposed to be in part due to inhibition of microglial activation^41^ as well as mitigation of myeloid cell entry into the brain thus reducing inflammation. Studies also suggest that minocycline influences neuroplasticity by promoting dendritic spine development and maturation,^42^ and neurite outgrowth.^43^

All rodent neurobehavioral testing was performed using standard protocols by the Emory University Rodent Behavioral Core (RBC). The rodent behavioral core was blinded to the disposition and genotype of the mice. Unless where explicitly stated, all data, including image analysis was performed by individuals who were blinded to the genotype and/or disposition of the mice. After neurobehavioral tests were completed, mice were sacrificed and the brain extracted for histological analysis. Immunohistochemical assay was performed to (1) ascertain/quantify the presence of inflammatory cells (activated microglia and bone marrow-derived microglial) and (2) quantify and phenotype proliferating cells in the dentate gyrus (i.e. cell fate). Golgi-Cox impregnation was used to determine the density of dendritic arbors and spines, as well as dendritic spine morphology. The assignment of mouse brain for either immunohistochemistry or Golgi-Cox impregnation was done randomly within each treatment group and genotype.

### Minocycline treatment

To determine whether neuroinflammation was a mechanism for cognitive and neurobehavioral deficits in aged sickle cell mice, we administered minocycline at a dose of 90mg/kg/day orally via drinking water for 3 weeks. The amount of water drank by the mice per cage was measured and the drug dosing adjusted every day based on this value. This was to ensure consistency in the dosage of drug that the mice were getting each day. Sickle cell mice acting as controls were given plain drinking water.

### Novel Object Recognition Test

This is a test of hippocampal-based learning and memory. It was adopted for our study because as in their human counterpart, sickle cell mice are also predisposed to developing pain crises, which could bias the results of a Morris water maze test for instance. This test is conducted in an open field made of transparent plexiglass and measuring 12” X 12” X 15” (L x W x H). Initially, animals were allowed 2 – 3 days of habituation which consisted of allowing them to explore the open field for 5 minutes each day. The total distance covered during this time as well as the time spent in the center of the open field was recorded and provides a measure of the animal’s behavior (anxiety/depression). On the 3^rd^ or 4^th^ day, the mice were trained by placing two identical objects (test objects) in the enclosure and allowing the mice to explore it for 5 minutes. Following training, the mice were returned to their home cages for about 30 minutes. After this delay, the learning and memory of the mice were tested by replacing one of the test objects used during training with a novel object in the open field. The time spent investigating the familiar (test object) and the novel object was recorded over a 5 minutes period. Mice were removed and returned to their home cages at the end of the test session. The enclosure as well as the test and novel objects, are wiped with alcohol between animals/tests.

### Fear Conditioning Test

This test measured the ability of mice to form and retain the memory of an association between an aversive experience and environmental cues. We used a standard fear conditioning paradigm, conducted over 3 days. On the first day, the mice were placed in the fear conditioning apparatus (7” W X 7” D X 12” H, Coulbourn Instruments, Holliston, MA) and allowed to explore for 3 minutes. After the period of habituation, we presented the mice with 3 pairs of conditioned stimulus (CS) – unconditioned stimulus (US) with a 1 minute inter trial interval between stimulus presentations. The CS was a 20 seconds long 85db tone and the US was 2 seconds of a 0.5 mA electric shock to the footpad/paw, timed to co-terminate with each CS presentation. It should be noted that this level of shock is relatively modest and very well tolerated by comparison to that in other studies such as seen in stress-induced drug reinstatement. The shock is delivered by a Precision Animal Shocker (Coulbourn Instruments, Holliston, MA) connected to each fear-conditioning chamber, with shock level set and verified before every training session. One minute after the presentation of the last CS-US presentation, the animals were returned back to their home cages. On day 2, the mice were presented with a context test, which involves placing them in the same chamber used for training on day 1, but this time, no shock is delivered. The amount of freezing per minute was recorded over 9 minutes or 540 seconds. On day 3, the mice were presented with a tone or cued test by exposing them to the CS in a novel environment/compartment. Initially the animals were allowed to explore the novel environment for 2 minutes. Following this brief period of habituation, the 85db tone was presented to the mice every minute for 9 minutes and the amount of freezing behavior per minute was recorded. In both the contextual and cued fear test, freezing behavior indicates a memory for either the context in which the shock was delivered or for pairing the tone with a shock. More freezing indicates better learning and thus more memory for the context and/or the CS-US pairing, while lesser freezing behavior indicates the opposite.

### Immunohistochemistry

At the conclusion of all the cognitive and neurobehavioral tests, animals were transferred to their home cages and were sacrificed the next day for brain extraction and histology. Briefly, animals were sacrificed using an overdose of pentobarbital and then perfused fixed with 4% (w/v) paraformaldehyde in PBS. Brains were extracted and post-fixed overnight at 4°C in 4% (w/v) PBS. In each mouse, the entire brain was sectioned at 50µm thickness. Using stereotactic coordinates,^44^ the sections that contain the hippocampus (dentate gyrus) were identified and used for immunohistochemistry in order to define the role of inflammation (neuroinflammation) on observed cognitive and/or neurobehavioral impairment in the sickle cell mice. We also attempted to map the predominant fate of new cells that were identified in the dentate gyrus. To enable us accomplish all these, we developed four antibody panels, thus, **panel 1** = anti-mouse Neun (AB90, Millipore, 1:500), anti-mouse Iba1 (019-19741, Wako Chemicals, 1:500), anti-mouse CD45 (ab23910, Abcam, 1:500); **panel 2** = anti-mouse BrdU (ab8152, Abcam, 1:500), anti-mouse Iba1 (019-19741, Wako Chemicals, 1:500), anti-mouse CD45 (ab23910, Abcam, 1:500); **panel 3** = anti-mouse BrdU (ab8152, Abcam, 1:500), anti-mouse doublecortin (DCX) (ab207175, Abcam, 1:500); **panel 4** = anti-mouse Neun (AB90, Millipore, 1:500), anti-mouse doublecortin (DCX) (ab20717, Abcam, 1:500), anti-mouse glial fibrillary acid protein (GFAP) (ab4674, Abcam, 1:1500) and starting with the first section showing the dentate gyrus to the last section, we labeled every 4^th^ brain slices with one of the antibody panels described above; i.e., slice 1 goes to panel 1, slice 2 to panel 2, slice 3 to panel 3 and slice 4 to panel 4, and then the process is repeated starting with panel 1. The general approach used for immunohistochemistry in our laboratory has already been published.^34^ For primary antibody labeling, free-floating tissue sections (50µm thick) obtained from post-fixed brain samples, were placed in a scintillating glass vial containing antibodies (diluted in antibody diluting solution for a total of 2mls volume) from one of the panels described above and after incubating for 16-24 hours at room temperature on a nutating shaker, the sections were washed with PBS and then incubated for 2 hours with the respective secondary antibody. Sections were mounted using Fluoromout G and imaged at 20X using a confocal microscope (Leica SP8, Leica Microsystems Inc.).

#### Immunohistochemistry and digital imaging analysis for Iba-1 and CD45

This assay was done to determine the presence of peripherally derived inflammatory (CD45^+^ or bone marrow-derived microglia) cells and/or activated microglia (Iba-1^+^) in and/or around (50 pixels on either side) of the dentate gyrus. The general approach for immunohistochemical labeling for Iba-1 and/or CD45 has been described above. For each mouse, images were taken using a 20X objective, from the dentate gyrus (DG) of the hippocampus. Image analysis was performed using NIH ImageJ software. For CD45, cells with positive labeling were counted in each dentate gyrus section. Analysis of Iba-1 labeling for activated microglia was done using a combination of digital image analysis -and visual confirmation of morphological features of activated microglia.^19, 22, 45^ In brief, a threshold for positive labeling was determined for each image that included all cell bodies and processes but excluded background staining. The number of CD45^+^ cells and activated microglia was then expressed as a density per 1000µm^2^ of dentate gyrus area.

#### 5’Bromodeoxyuridine (BrdU) and doublecortin (DCX) labeling

We examined the proliferation of actively dividing cells in the dentate gyrus (DG) using 5’bromodeoxyuridine (BrdU) labeling. BrdU (10 mg/ml; Sigma-Aldrich) was dissolved in warm PBS and then filtered. On the last 3 days prior to cognitive and neurobehavioral testing, mice were injected intraperitoneally with 50 mg/kg BrdU at 9:00 A.M. Brains were collected for BrdU immunohistochemistry. For quantification of BrdU positive (BrdU^+^) and doublecortin-positive (DCX^+^) cells, every fourth section throughout the hippocampus/dentate gyrus was collected and stained with **panel 3** = anti-mouse BrdU (ab8152, Abcam, 1:500), anti-mouse doublecortin (DCX) (ab207175, Abcam, 1:500) as described earlier. Sections were mounted on slides, coverslipped with Fluoromount G (Beckman Coulter), and stored at 4°C. Fluorescent images were captured using a 20X objective, on a confocal microscope (Leica SP8, Leica Microsystems Inc.). The image analysis was done using NIH ImageJ software for the total number of BrdU^+^ cells, DCX^+^ cells and BrdU^+^/DCX^+^ in each dentate gyrus section. As was the case with Iba-1 and CD45 positive cell quantification, the total number of BrdU^+^ cells, DCX^+^ cells and BrdU^+^/DCX^+^ in the DG were normalized per 1000µm^2^ of dentate gyrus area. This experiment enabled us to quantify the population of the proliferating cells (BrdU^+^ cells) that are neural progenitor cells (DCX^+^).

#### Phenotyping BrdU^+^ cells

We then proceeded to determine the phenotype of proliferating (BrdU^+^) cells in the dentate and/or “peri-dentate” gyrus area. Thus, brain slices were obtained as described earlier and triple labeled as follows; **panel 2** = anti-mouse BrdU (ab8152, Abcam, 1:500), anti-mouse Iba1 (019-19741, Wako Chemicals, 1:500) and anti-mouse CD45 (ab23910, Abcam, 1:500); **panel 3** = anti-mouse BrdU (ab8152, Abcam, 1:500), anti-mouse doublecortin (DCX) (ab207175, Abcam, 1:500); **panel 4** = anti-mouse NeuN^+^ (AB90, Millipore, 1:500), anti-mouse doublecortin (DCX) (ab20717, Abcam, 1:500), anti-mouse glial fibrillary acid protein (GFAP) (ab4674, Abcam, 1:1500). Panel 2 enables us to determine whether the proliferating cells are from the peripheral immune system or dividing microglia, panel 4 allows us to determine whether neural progenitor cells (NPCs) (expressing DCX) are more likely young or mature neurons (NeuN^+^) vs. astrocytes (GFAP^+^). Fluorescent images were captured using a 20X objective on a confocal microscope (Leica SP8, Leica Microsystems Inc.). The image analysis was performed using NIH ImageJ software for the total number of DCX^+^/NeuN^+^ cells or DCX^+^/GFAP^+^ cells in each dentate gyrus section. As was the case with Iba-1 and CD45 positive cell quantification, the total number of DCX^+^/NeuN^+^ cells or DCX^+^/GFAP^+^ in the DG were normalized per 1000µm^2^ of dentate gyrus area. A previous study was used as a reference for morphological comparison of immature and mature DCX^+^ neurons.^46^

#### Quantification of dendrites arbors, dendritic spine density and spine morphology

This experiment was performed with the Golgi-Cox impregnation approach, using the FD Rapid GolgiStain™ Kit according to the manufacturer’s instructions. ^25, 26, 28, 47-49^ It allowed us to visualize and characterize the remodeling/plasticity of pyramidal neurons in the cortical and hippocampal/dentate gyrus regions of the brain. Brightfield images were acquired using a 60X using an oil immersion objective mounted on a fluorescent microscope.

The detailed approach for Golgi-Cox impregnation has already been published. But Briefly, mice were sacrificed using an overdose of pentobarbital (150 mg/kg, intraperitoneally), and their brains were removed, rinsed with Nanopure water, and immersed in the impregnation solution composed of potassium dichromate, mercuric chloride and potassium chromate. The brains were stored at room temperature for 2 weeks and then transferred and stored in the cryoprotectant solution for 2.5 days in the dark. Coronal sections of 100 – 200 μm thickness were cut on a vibratome with the tissue bath filled with the cryoprotectant solution. Sections were mounted on gelatin-coated glass slides (Electron Microscopy Solution Inc., Hatfield, PA), and allowed to air dry at room temperature in the dark for 2 – 3 days. After drying, the sections were rinsed with Nanopure water, reacted in the working solution, and dehydrated with a 50, 75, 95 and 100% graded ethanol series. Finally, the sections were defatted in xylene and cover-slipped using Permount (VWR International LLC., Radnor, PA).

Digitized images were acquired at using a 60X (oil immersion) objective on an upright fluorescent microscope. Z-stacks of pyramidal neurons were obtained from the hippocampus/dentate gyrus and the cerebral cortex in a clockwise fashion to ensure reproducibility. We quantified the dendritic spine density and morphology using manual visual rating and published criteria, with the rater blinded to the genotype and disposition of the animals.^50-52^ The z-stacks were then projected using minimum intensity projection and the dendrite arborization pattern and thus branch number/density was quantified using an adaptation of the Sholl analysis (http://imagej.net/Sholl_Analysis).^53-56^ Results of dendritic spine quantification were expressed per 10 µm dendrite segment.

### Statistical consideration

A 2-way Analysis of Variance (ANOVA) was used to compare groups on performance in respective neurobehavioral and cognitive tests that have a time component (Open field, distance traveled and fear conditioning). A one-way ANOVA was used to compare differences in percent preference for NOR and density of the different cells quantified in the dentate or “peri-dentate” gyrus area. Also, a one-way ANOVA was used to compare differences in dendrite density and dendritic spine density as well as proportion of immature dendritic spines. When the data is not normally distributed, a non-parametric ANOVA was used instead. A p-value of <0.05 was used as indicating a statistically significant difference between genotype groups and/or treatment groups. Unless otherwise stated, absolute values are presented as mean ± SE.

## Results

### Aged sickle cell mice displayed significant cognitive and neurobehavioral deficits compared to control mice

For this study, we examined the learning, memory and anxiety-like/depression behaviors in 13 months old Townes sickle cell and control mice using a NOR test. Sickle cell mice exhibited evidence of significant anxiety-like behaviors and depression indicated by the shorter distance traveled (*p=0.0004*, Fig 1i) and thigmotaxis (p=*0.002;* Fig 1ii) when compared to controls. Sickle cell mice spent significantly shorter investigation time (2.39 sec ± 0.64) on the novel object, compared to control (8.62 sec ±1.80; p=*0.007*; Fig 1iii). Similarly, sickle cell mice showed learning and memory deficits indicated by a lower percent preference for the novel object (32.15%± 7.8), compared to controls (58.3% ± 9.9; p=*0.010*; Figure 1iv).

**Figure 1.**
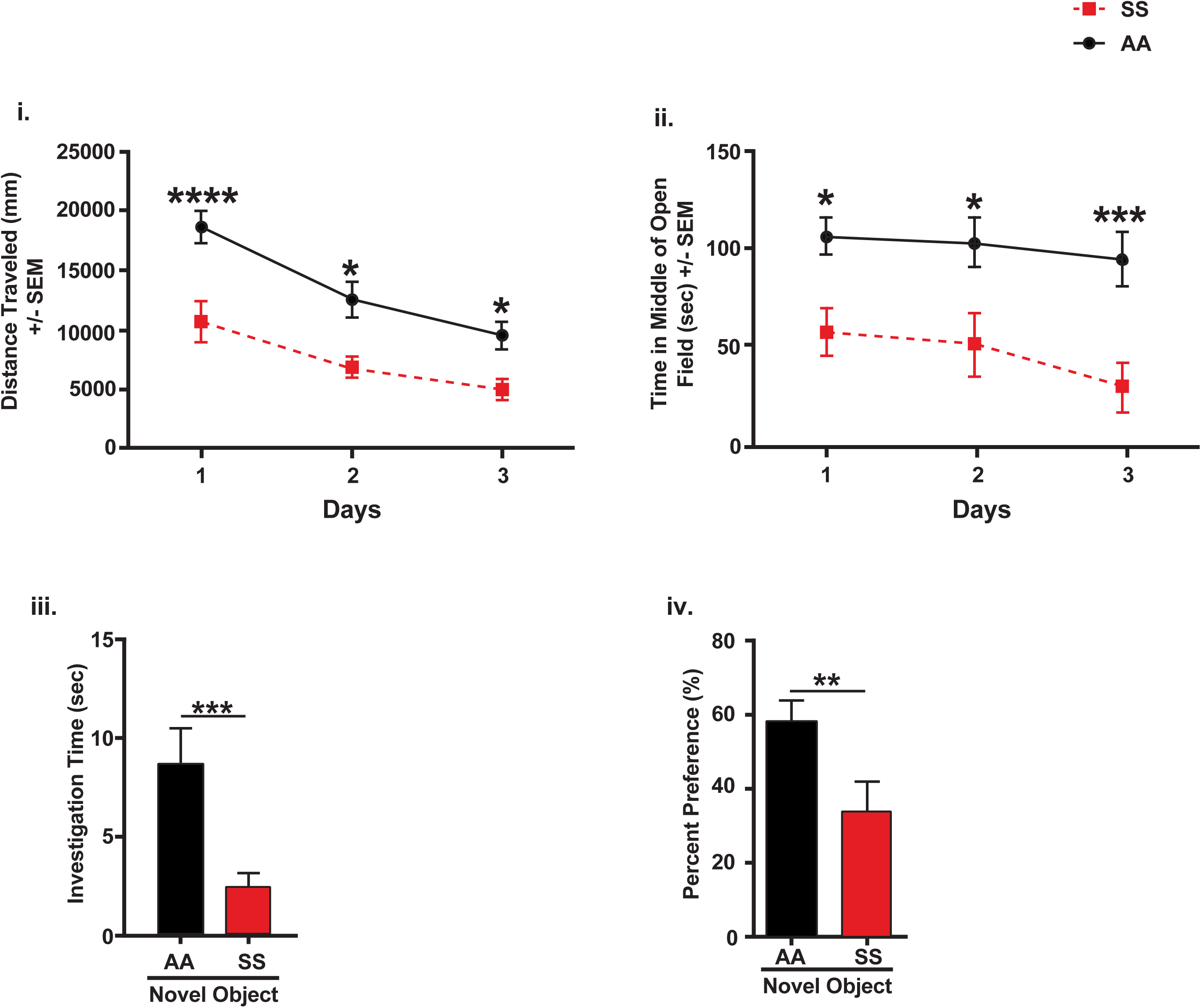
Abnormal cognition and behavior in 13 months old sickle cell compared to control mice. In the figure, (i) distance traveled (*p=0.0004*), (ii) time spent in the open field *(p=0.002)*, (iii) investigation time on the novel object (p=*0.007;* and (iv) percent preference for the novel object *(p=0.01)* were significantly lower in sickle cell mice (SS) compared to controls (AA) indicating that aged (13 months old) sickle cell mice have cognitive and neurobehavioral abnormalities compared to controls. Data is presented as mean ± SEM. N = 10 per group

### Aged sickle cell mice displayed cognitive and neurobehavioral deficits that were not seen in younger sickle cell mice

To demonstrate this, we examined the learning, memory and anxiety/depression behaviors in 6 months old Townes sickle cell and control mice using a novel object recognition and open field test as was done for the original cohort of 13 months old sickle cell and control mice. There were no significant deferences in neurobehavioral characteristics between sickle cell and controls mice as evidenced by similar measures for distance traveled (*p=0.85;* Fig 2i) and thigmotaxis (*p=0.55;* Fig 2ii). There was also no significant difference in learning and memory between sickle cell and control mice at 6 months as they showed similar investigation time (2.64 sec ± 1.20 vs. 2.34 sec ± 0.72; *p=0.25*; Fig 2iii) on the novel object as well as percent preference for the novel object (28.39% vs 45.92%; *p=0.21;* Fig 2iv). The difference in behavior and cognition between sickle cell and control mice reappeared when the same mice were allowed to age to 13 months. The aged sickle cell mice exhibited more anxiety/depression-like behaviors indicated by significantly shorter distance traveled, (Fig 3ai) and thigmotaxis (Fig 3aii) and were less likely to spend time investigating the novel object. Aged sickle cell mice also had significantly lower percent preference or preference ratio for the novel compared to aged control mice (Fig 3aiv).

**Figure 2.**
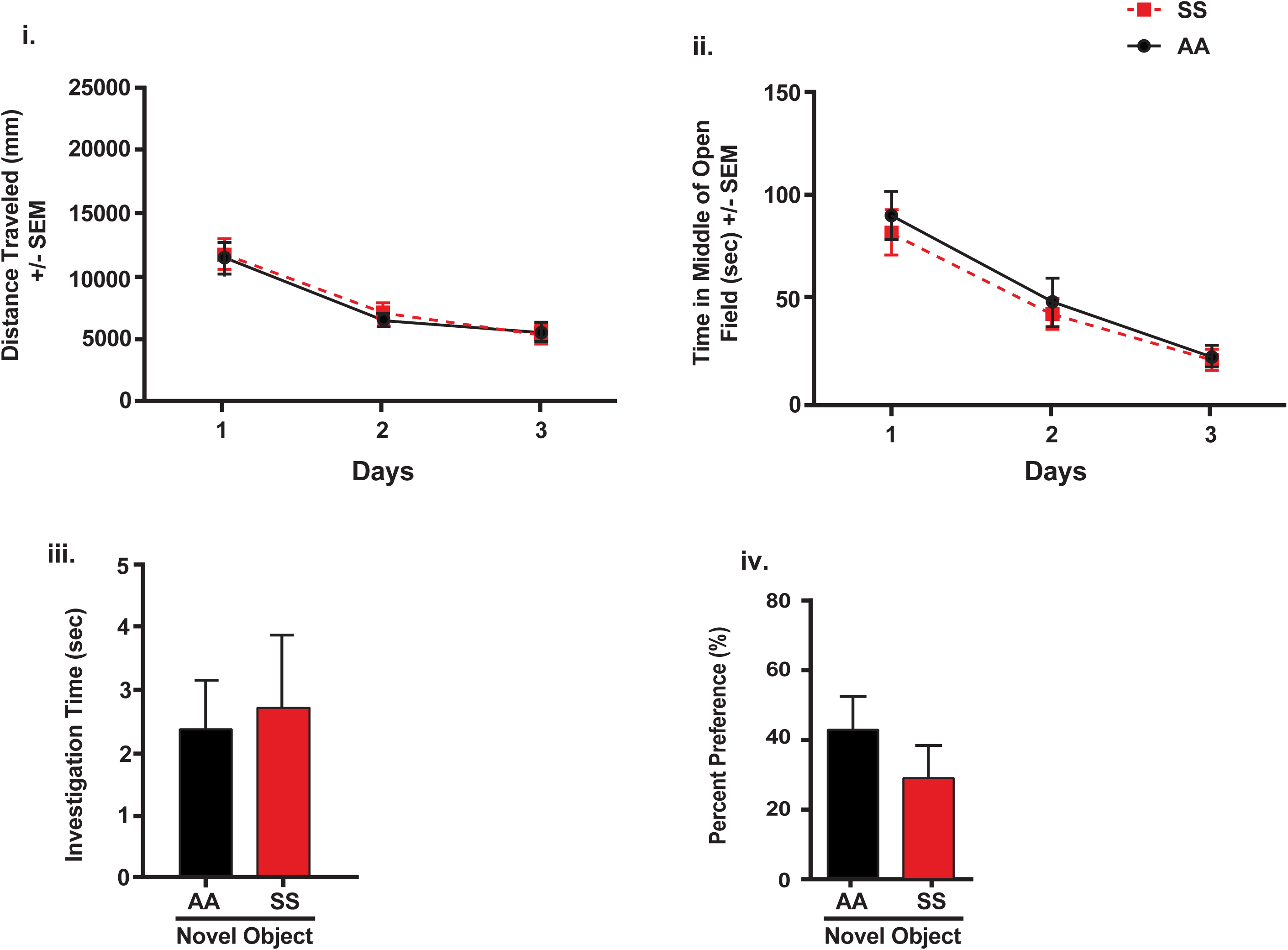
Similar levels of cognitive and neurobehavioral function in sickle cell compared to control mice. No significant cognitive or neurobehavioral abnormalities were seen in 6 months old sickle cell (SS) mice compared to controls (AA) as indicated by similar levels of (i) distance traveled, (ii) time spent in the open field (sec), (iii) time in second spent investigating the novel object and (iv) percent preference for the novel object. Result is presented in mean ± SEM. N = 15 sickle cell mice and 12 AA or control mice.

**Figure 3.**
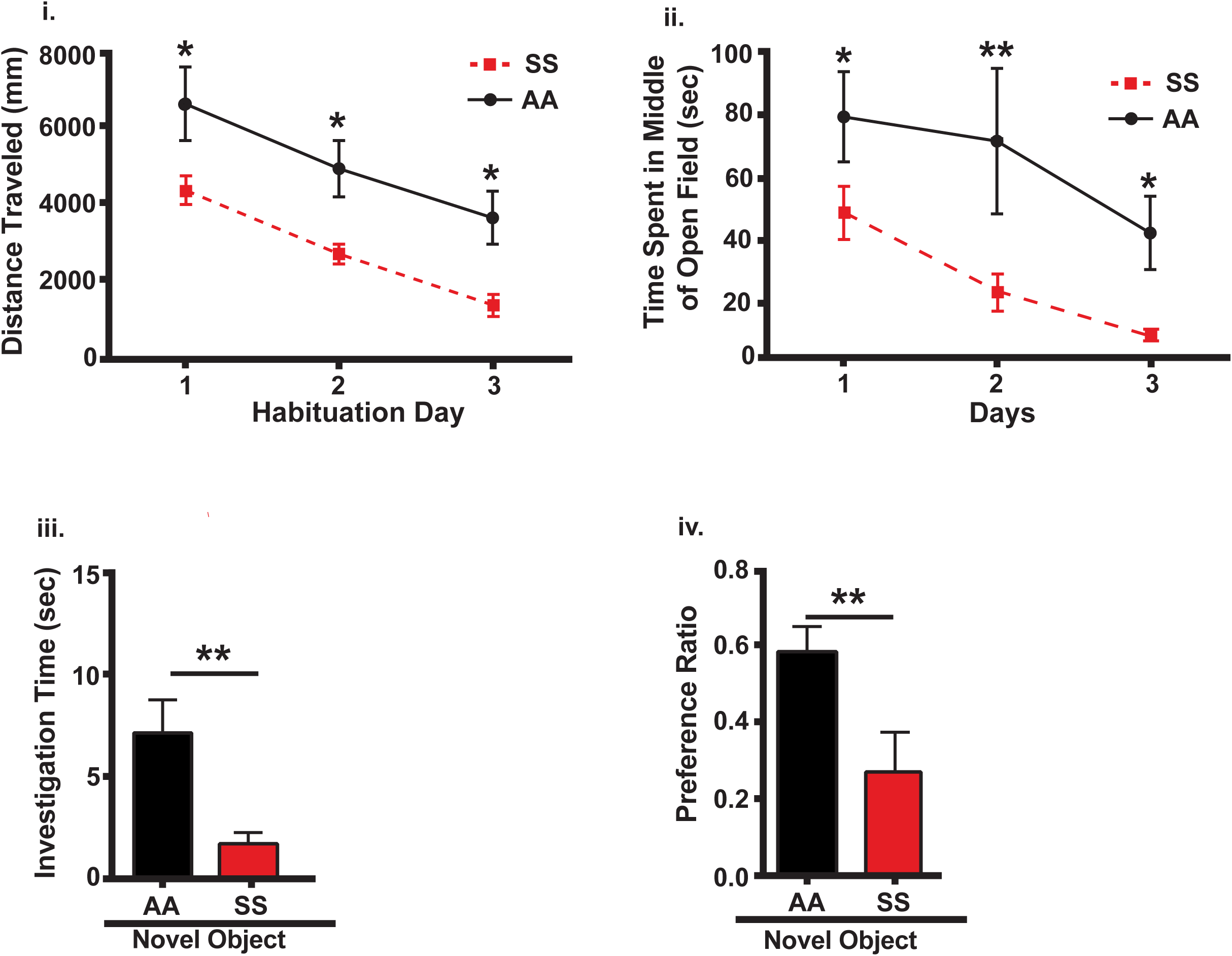
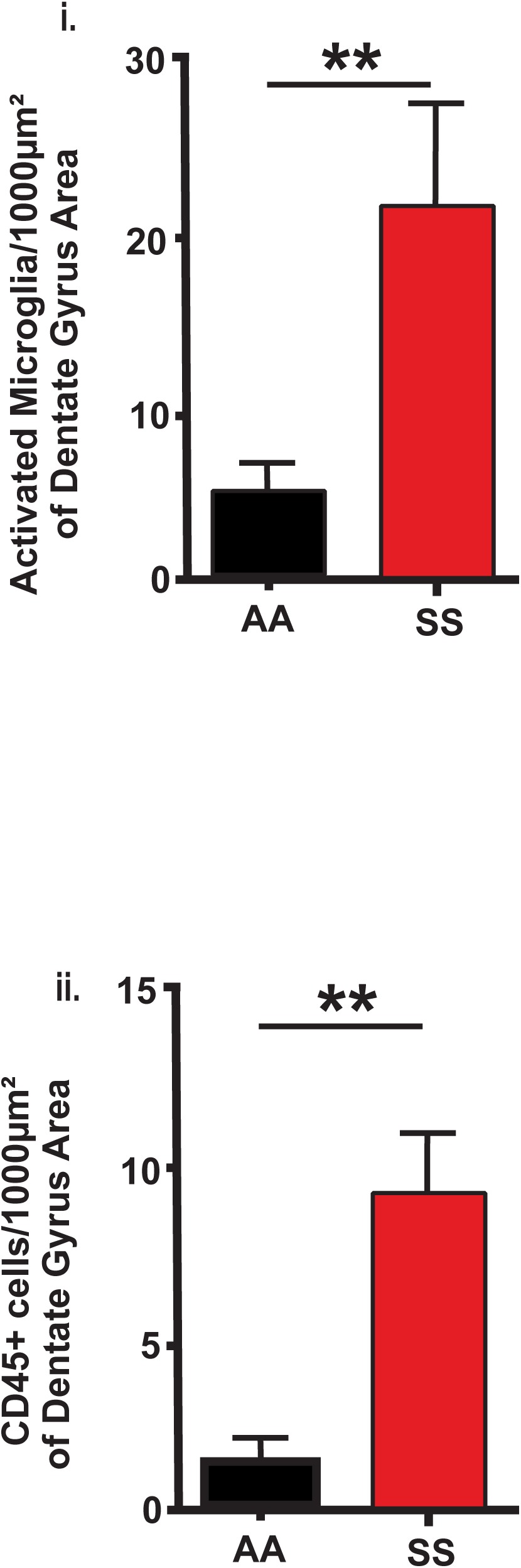
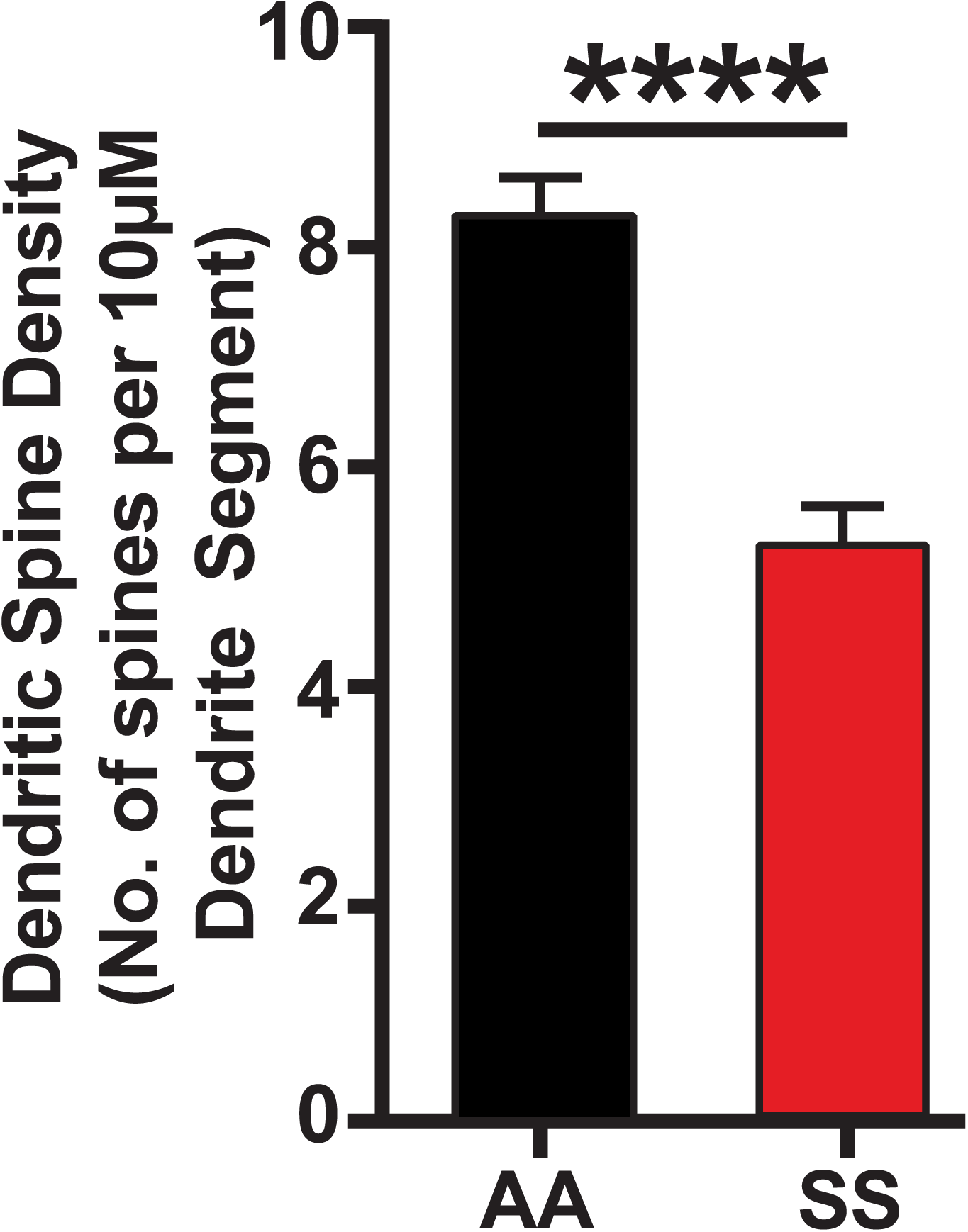
Cognitive and neurobehavioral abnormalities reappeared when young (6 months old) sickle cell mice (from Fig. 2) were aged to 13 months of age, compared to control mice. Aged sickle cell mice also had significant cellular evidence of neuroinflammation as well as lower dendritic spine density compared with controls. When aged to 13 months old, sickle cell mice (SS) mice had significantly more cognitive and neurobehavioral abnormalities that were not seen at 6 months of age when compared to controls (AA). The (i) distance traveled (*p=0.0008*; (ii) time spent in the open field (p=*0.0007*; (iii) investigation time on the novel object (*p=0.02*) and (iv) percent preference for novel object where significantly lower when compared to AA mice (*p=0.02*). In **Figure 3b**, we show higher levels of neuroinflammation (i.e more pro neuroinflammatory cells) in sickle cell mice when compared to controls. At baseline, there were more (i) activated microglia (*p=0.005)*, measured via Iba1 labeling and (ii) peripherally (bone marrow) derived mononuclear cells (*p=0.001*), measured via CD45 labeling, in the dentate and prei-dentate gyrus of area of sickle cell mice compared to controls. In **Figure 3c**, Dendritic spine density was significantly reduced in sickle cell mice compared to controls. Aged (13 months old) SS mice had significantly lower dendritic spine density compared to AA mice (*p=0.0002).* Results are presented as mean ± SEM. N = 9 controls or AA mice, 8 SS treated mice and 6 SS non-treated mice.

### Aged sickle cell mice have more evidence of neuroinflammation compared to controls

Immunohistochemical analysis of brains from the 13 months old mice above shows that sickle cell mice had significantly more evidence of neuroinflammation than controls, indicated by a significantly higher number of activated microglia per 1000μm^2^ of dentate gyrus area (22.01 ± 6.17 vs. 6.02 ± 1.44; *p=0.005*; Figure 3bi). SS mice also had a significantly higher number (9.32 ± 2.23) of CD45^+^ peripherally derived mononuclear (or bone marrow-derived microglia) cells per 1000μm^2^ of dentate gyrus area compared to controls (1.57 ± 0.56; *p=0.001;* Figure 3bii).

### Aged sickle cell mice have lower dendritic spine density compared to controls

We assessed dendritic spine density in a subset of 13 months old sickle cell and control mice in order to determine any underlying neuronal changes that could be responsible for the observed cognitive and neurobehavioral deficits. As shown in Fig. 3c, we observed a significantly lower dendritic spine density per 10µm of dendrite segment in sickle cell mice (5.62 ± 0.45) compared to control mice (8.46 ± 0.37; *p=0.0002).*

### Treatment with minocycline restored cognitive and neurobehavioral deficit in aged sickle mice

Next we randomized half of the 13 months old sickle cell mice from the experiment in Fig. 2 to either receive minocycline (90mg/kg) in their drinking water or plain drinking water. We had performed a pilot experiment to ensure that minocycline had no impact on the amount of water drank (data not shown), since better hydration can improve sickle cell outcomes in general. Cognitive function (learning and memory) and behavior in sickle cell mice that received minocycline improved to the same levels as non-sickle cell mice controls. On the other hand, cognitive function and behavior among sickle cell mice that did not receive minocycline were little changed or slightly worsened. The result for this specific cohort is presented in the Online-only supplementary results. We repeated this experiment in a larger of cohort of aged (13 months old) mice (n = 15 – 20 mice per group) with an addition of 5 non-sickle cell (AA) mice group to receiving minocycline (result not included). There was no significant difference between treated sickle cell and treated control mice and thus we have decided not to include treated controls in the presentation of our results. Our results from the open field test showed that treated sickle cell mice had reduced evidence of anxiety/depression similar to control levels, but better than non-treated sickle cell mice, indicated by a significantly longer distance traveled (*p=0.02*; Fig 4ai) reduced thigmotaxis by spending more time in the middle of the enclosure (*p=0.04*; Fig 4aii). Similarly, learning and memory for treated sickle cell mice measured as percent preference for the novel object (72.21% ± 8.92) were the same as that of controls, but significantly better than non-treated sickle cell mice (14.86% ± 5.98; *p*<*0.0001*; Fig. 4aiii).

**Figure 4.**
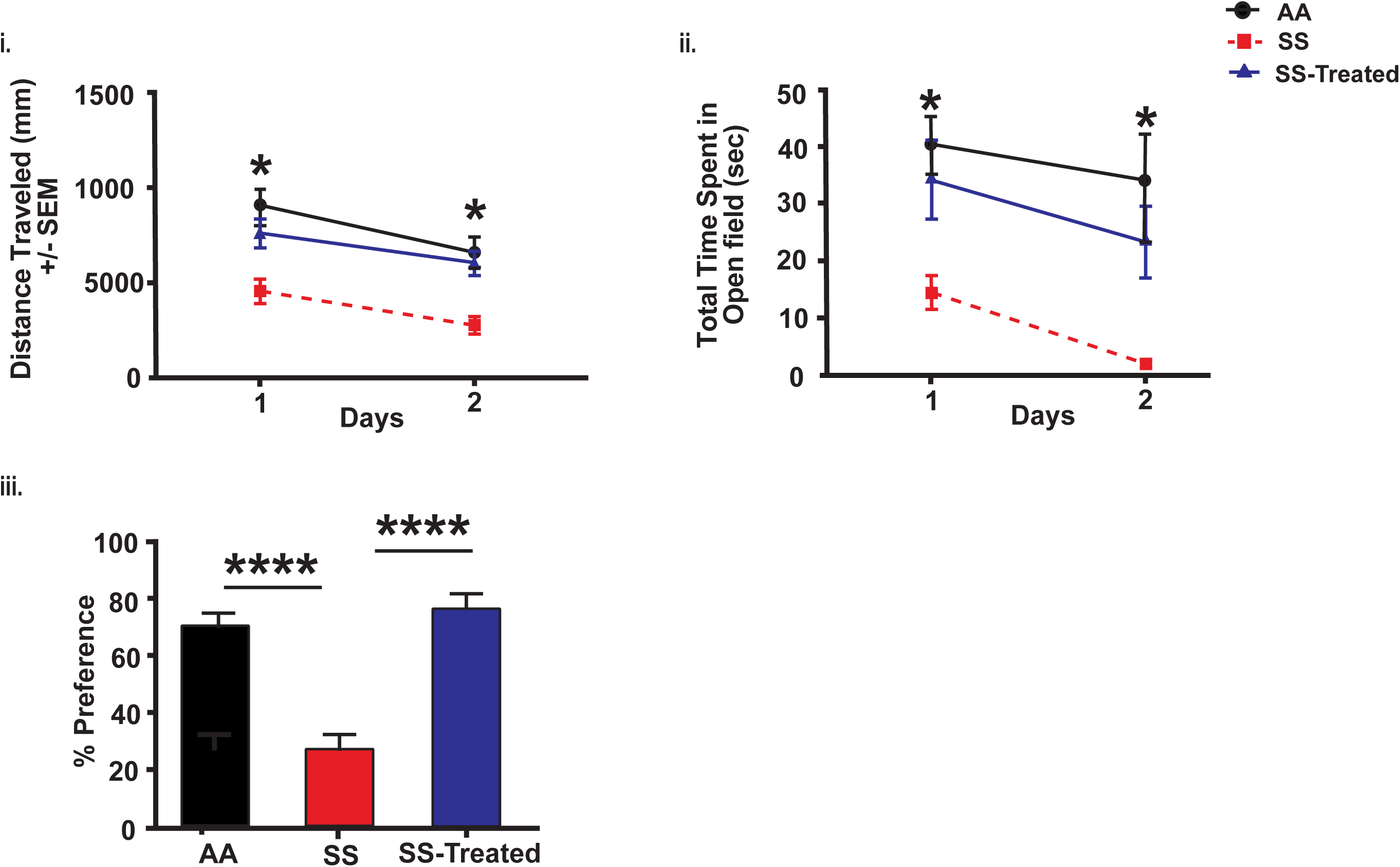
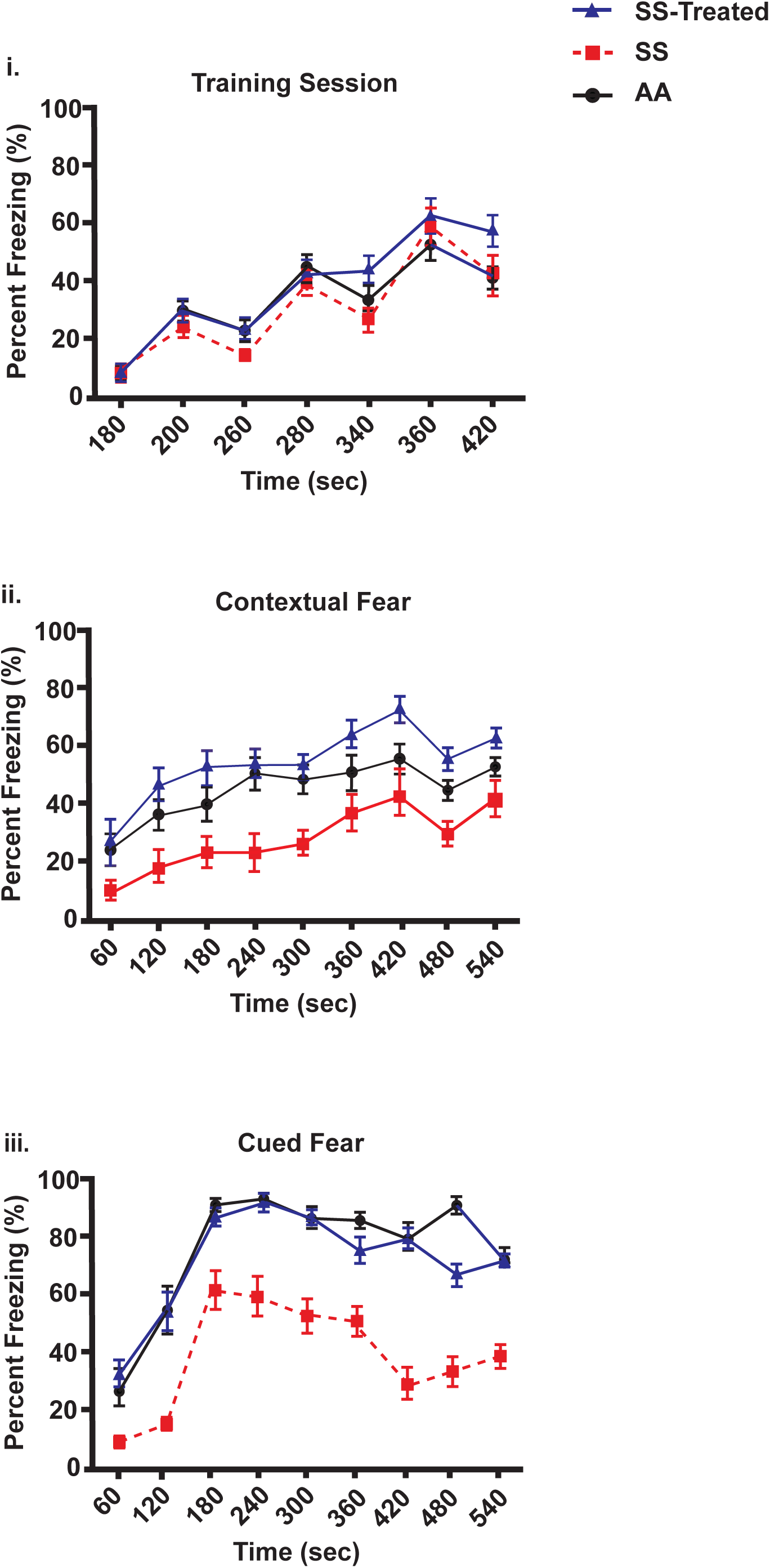
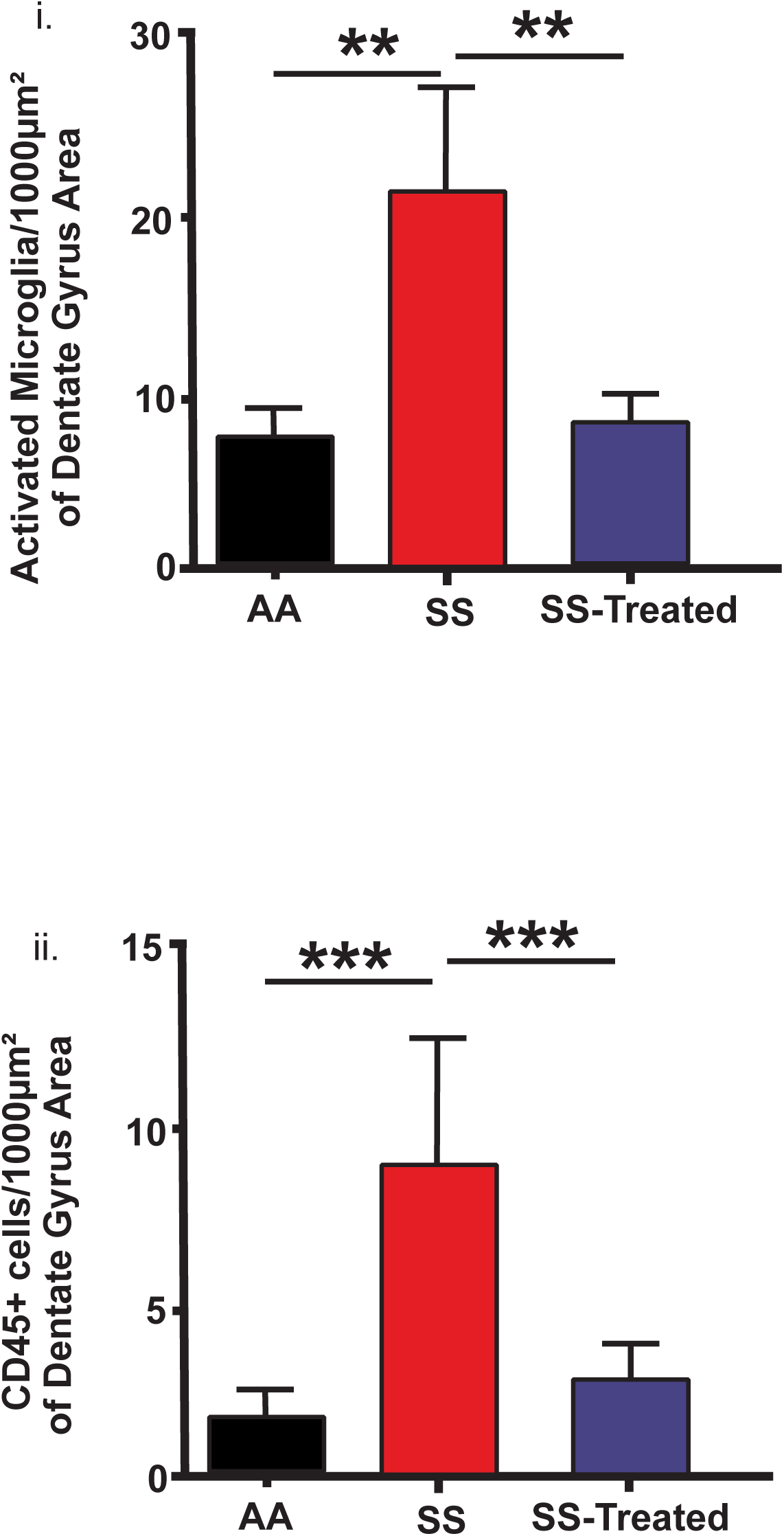
Minocycline treatment attenuates cognitive and neurobehavioral abnormalities and evidence of neuroinflammation in sickle cell mice compared to non-treated sickle cell mice. **Figure 4a** shows there was significant improvement in (i) Distance traveled (*p*<*0.0001)*, (ii) time spent in open field (p=*0.008*) and percent preference for the novel object (*p*<*0.0001*) in treated sickle cell mice (SS-treated) compared to non-treated sickle cell mice (SS), non-treated sickle cell mice were not diferent from data observed in our prior experiments in Figs. 1 and 3a. Cognitive and neurobehavioral performance were similar between SS-treated mice and controls. In **Figure 4b**, fear conditioning test shows that treatment with minocycline attenuates congnitive and neurobehavioral deficits in treated sickle cell mice compared to non-treated sickle cell mice. The figure shows (i) comparable levels of performance on the acquisition phase of fear training, evidenced by similar percent freezing among non-treated sickle cell (SS), treated sickle cell (SS-treated) and control (AA) mice (*p=0.22*). In the testing phase, treated sickle cell mice performend significantly better on the (ii) contextual fear testing with significantly higher percent freezing at all time points compaired to non-treated sickle cell mice (*p*<*0.001*). Similarly, on (iii) cued fear test, sickle cell mice treated with minocycline also had significantly higher percent freezing at all time points compared with non-treated sickle cell mice (*p*<*0.001*). Comparison of percent freezing between controls mice and non-treated sickle cell mice was also statisitically singficant, while there was no significant difference in percent freezing between treated sickle cell mice and controls. It is noted that higher percent freezing indicates bettwe memory for the conditioned and unconditioned stimulus and thus better learning, memory and thus cognitive function. In **Figure 4c** we show that minocycline treatment attenuates levels of neuroinflammation in sickle cell mice when compared to non-treated sickle cell mice. Significantly fewer numbers of (i) activated microglia (*p=0.007*), measured via number of labeled Iba1 cells, and peripherally derived mononuclear cells (*p=0.002*), measured via number of labeled CD45 cells, was observed in treated sickle cell mice (SS-treated) when compared to sickle cell mice (SS). Results are presented here as mean ± SEM. N = 15 mice per group for cognitive and neurobehavioral testing (**Figs. 4a** and **4b**) and 9 per group for histological analysis.

We further confirmed impaired cognitive (learning and memory) and neurobehavioral function as well as improvement with minocycline treatment using the fear conditioning test. In the acquisition phase, there were no significant differences between sickle cell mice, controls or sickle cell mice on minocycline treatment. The percent freezing was 7.6%, 7.7% and 10.2%, respectively in sickle cell mice, control mice and sickle cell mice on minocycline treatment, at the acquisition phase (Fig. 4bi). Thus percent freezing during the acquisition phase was essentially similar between all genotype and treatment groups except at the 340 secs (*p=0.008*) and 420 secs ((*p=0.02*) time points where treated sickle cell mice or controls seemed to do better than non-treated sickle cell mice. On contextual fear testing, sickle cell mice treated with minocycline had significantly higher freezing percent across all time points, an indication of better learning and memory compared to sickle mice that were not treated with minocycline or that received plain drinking water. There was no significant difference in performance on this test between sickle cell mice and controls except at the 360 and 420 seconds time point (Fig. 4bii). Similarly, on cued fear testing, sickle cell mice that received minocycline also showed significantly better memory for the conditioned stimulus at all time points, compared to sickle cell mice that received plain drinking water. And as was the case with contextual fear testing, there were no significant differences between treated sickle cell mice and controls except at the 480 seconds time point (Fig. 4biii).

### Treatment with Minocycline Attenuates Neuroinflammation in Sickle Cell Mice

Consistent with previous results, the treatment of sickle cell mice with minocycline resulted in a significant reduction in evidence of neuroinflammation. Thus, sickle cell mice that received minocycline showed significant reduction in the number of activated microglia (8.21 ± 1.93 cells/1000µm^2^ vs 22.01 ± 6.12 cells/1000µm^2^; *p=0.007*) and CD45^+^ bone marrow-derived microglia or mononuclear cells (3.04 ± 0.80 cells/1000µm^2^ vs. 9.32 ± 2.23 cells/1000µm^2^; *p=0.002*) compared to non-treated sickle cell mice. There was no statistically significant difference between treated sickle cell mice and controls, in the number of activated microglia or CD45^+^ bone marrow-derived microglia in the dentate or peri-dentate gyrus area. However, similar to the observation between treated and non-treated sickle cell mice, the number of activated microglia (*p=0.010*) and CD45^+^ bone marrow-derived microglia or mononuclear cells (*p=0.0006*) was significantly different between controls and non-treated sickle cell mice (Fig 4c i and ii).

### Sickle Cell Mice treated with minocycline showed improvement in neuroplasticity and potentially neurogenesis

Finally, we sought to determine the structural neuronal mechanism for the observed cognitive and neurobehavioral benefit of minocycline. We observed that sickle cell mice that received minocycline had significantly higher dendritic spine density (9.31 ± 0.69 spines/10µm vs. 5.95 ± 0.74 spines/10µm; *p*<*0.0001*, Fig. 5ai) compared to age-matched sickle cell mice that did not receive minocycline treatment or received plain drinking water. Consistent with previous results (Figure 3c), sickle cell mice that did not receive minocycline also had significantly fewer dendritic spines per 10µm of dendrite segment (5.95 ± 0.74 vs. 9.85 ± 1.87, *p=0.0001*; Fig. 5ai) compared to controls. With regards to spine morphology, we also observed that sickle cell mice treated with minocycline had significantly (more than 2-fold) fewer proportion (%) of immature dendritic spines as a percentage of the total spines counted (12.08 ± 1.57% vs. 32.08 ± 4.81%, *p*<*0.0001;* Fig. 5aii) compared to age-matched sickle cell mice that received plain drinking water. Similarly, non-treated sickle cell mice also had a more than 2-fold higher proportion of immature dendritic spines (32.08 ± 4.81 vs. 14.34 ± 2.48, *p=0.0002*; Fig. 5aii) when compared to controls. There was no significant difference in the proportion of immature spines between treated sickle cell mice and controls. In examining the impact of minocycline on neuroplasticity, we evaluated dendrite arborization using Sholl analysis. Compared to controls, non-treated sickle cell mice had significantly fewer dendrite arbors at 5μm (3.94 vs 5.33; *p=0.004)* and 10μm (5.69 vs 7.50; *p=0.005)* distance away from the soma (Fig. 5b i,ii, and iv). Also, non-treated sickle cell mice also had significantly fewer dendrite arbors compared to sickle cell mice that received minocycline. This difference was present at all distances from the soma that was evaluated and p-value at each distance was <0.01 (Fig. 5b i, iii and iv). These experiments show that the treatment of sickle cell mice with minocycline resulted in better neuroplasticity and our prior data above shows that this is likely mechanistically linked to improvement in cognitive and neurobehavioral functions.

**Figure 5.**
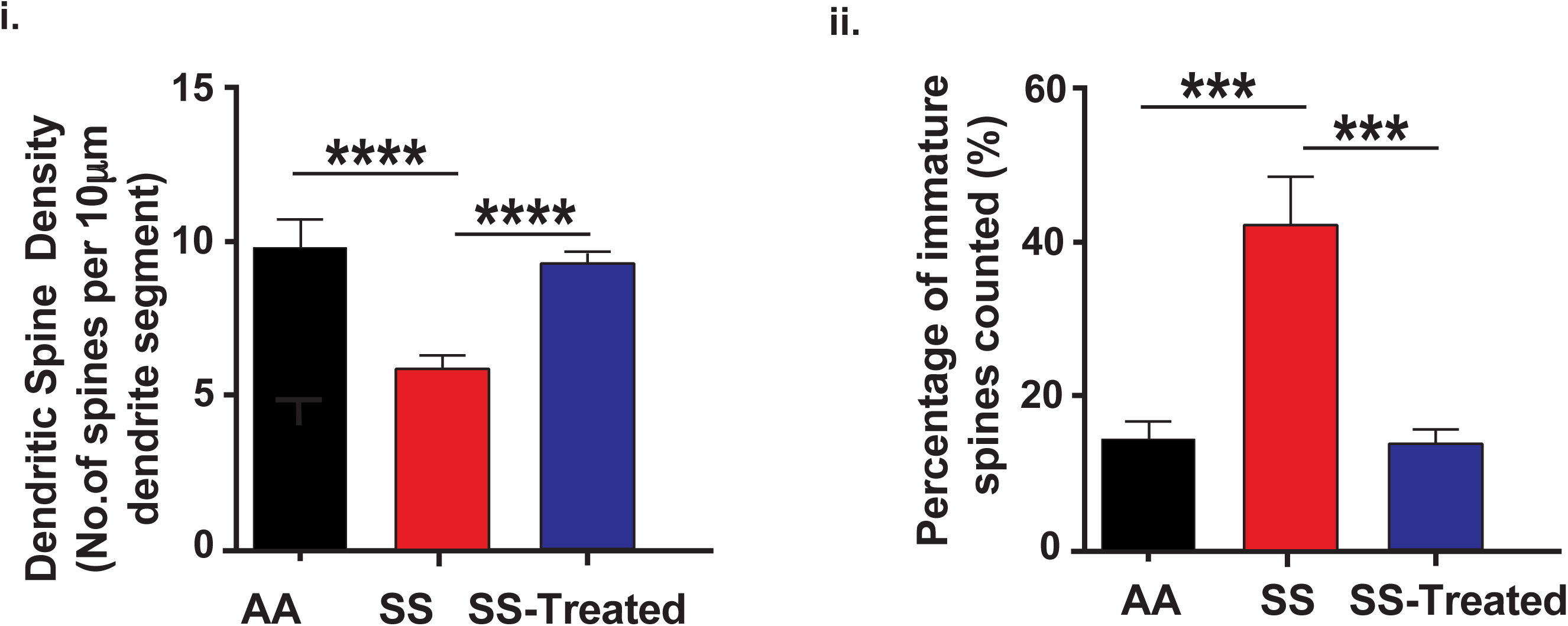
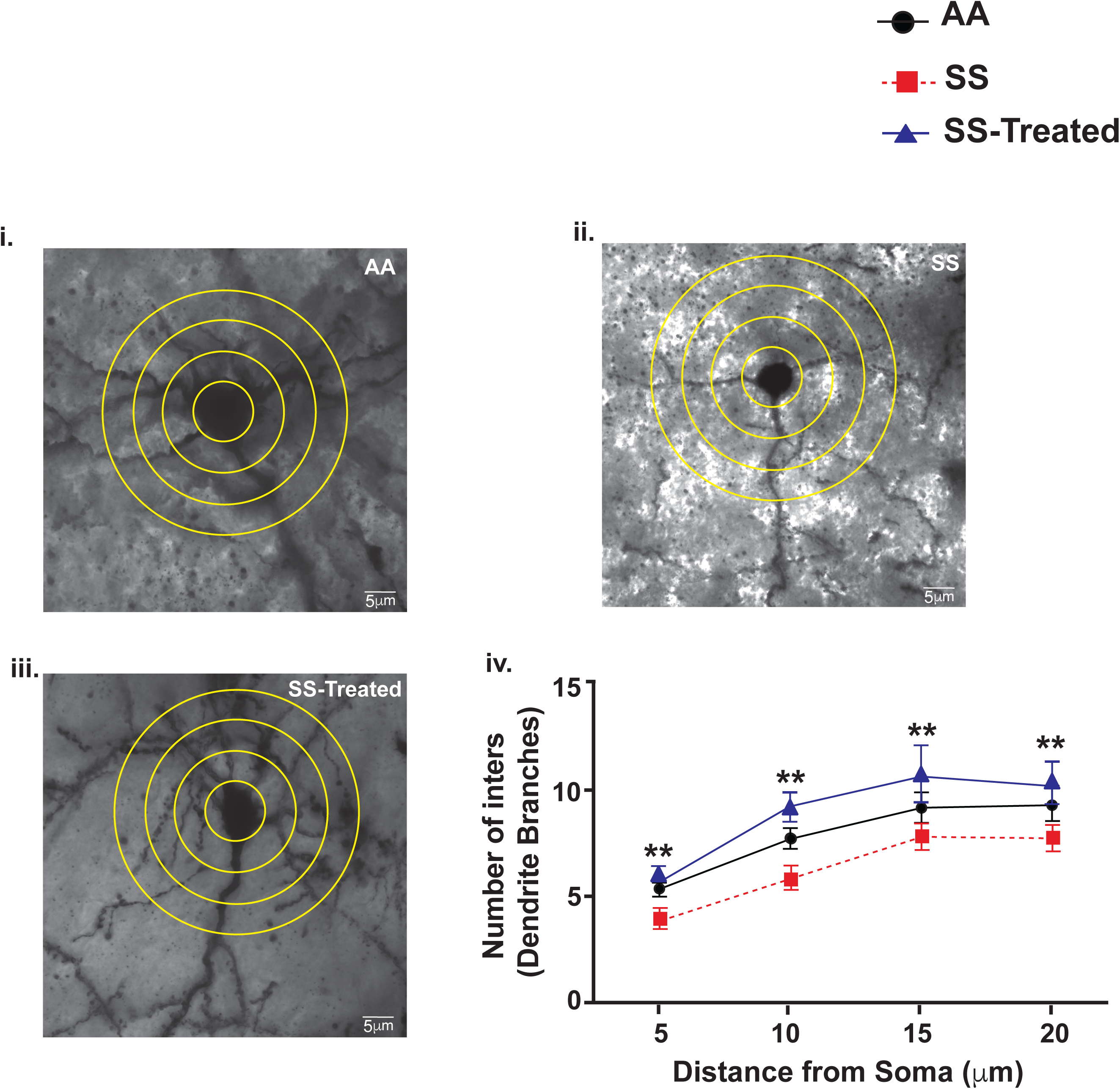
Minocycline treatment results in improvement in neuroplasticity evidenced by increased dendritic spine density and dendrite arbors, but a decreased proportion of immature spines in sickle cell mice treated with minocycline compared to non-treated sickle cell mice. In the figure 5a, we see that (i) Sickle cell mice treated (SS-treated) with minocycline had a significantly higher dendritic spine density (*p*<*0.0001*) per 10µm dendrite segment compared to non-treated sickle cell mice (SS). There were no significant differences in dendritic spine density between control (AA) and SS-treated mice. Also, sickle cell mice that did not receive minocycline (non-treated sickle cell mice or SS) had a 4-fold higher proportion of immature dendritic spines compared to sickle cell mice that received or were treated with minocycline i.e. SS-treated (*p*<*0.0001*) and controls (*p=0.0002*). In Figure 5b, we used Sholl analysis to evaluate the number of dendritic branches with 5µm interval between concentric circles (20μm distance total). Representative images are shown for (i) controls (AA), (ii) non-treated sickle cell (SS), and (iii) treated sickle cell mice (SS-treated). Figure 5biv shows that non-treated sickle cell (SS) mice had significantly fewer dendrite arbors compared to controls at 5µm (*p=0.004)* and 10µm (*p=0.005)* distance away from the soma. Similarly, SS mice had fewers dendrite arbors compared with treated sickle cell (SS-treated) mice at all distance from from the soma, i.e. at 5µm (*p*<*0.0001)*, 10µm (*p=0.0003)*, 15µm (*p=0.0487)*, and 20µm (*p=0.0280)* away from the soma. There was no significant difference seen between controls and treated sickle cell mice. The interval distance as well as the final distance of 20µm were chosen appriori based on the size of the image that could be generated from our fluorescent microscope. Results are presented as mean ± SEM. N = 6 per group. Note that these are the rest of the mice from (**Figs. 4a** and **4b**).

We sought to determine whether minocycline has an impact on cell fate for neuron progenitor cells (NPCs), due to its reported impact on inflammation and “spinegenesis”. Our final sample size was too small for a robust assessment of neurogenesis and as such the analysis did not shows statistically significant results (see Online only supplementary results). However, we observed a trend where sickle cell mice treated with minocycline had a higher number of dividing (BRDU^+^) cells when compared to non-treated sickle cell mice (Fig. S2a). Also, while treated and non-treated sickle cell mice showed similar numbers of DCX^+^ (NPCs) and BRDU^+^DCX^+^ cells (Fig. S2a). There seem to be a marked increased in the number of NPCs differentiating into mature neurons (DCX^+^NeuN^+^) cells among sickle cell mice treated with minocycline, while among sickle cell mice cells that did not receive minocycline, the NPCs seems to differentiate more into astrocytes (DCX^+^GFAP^+^) cells (Fig. S2b).

## Discussion

In this study, we showed that aged (13 months old) Townes sickle cell mice have more severe cognitive and neurobehavioral deficits compared to age and sex-matched controls, similar to what has been described in humans with SCD.^14^ There was also a significantly higher levels of neuroinflammation (evidenced by a higher density of activated microglia and bone marrow-derived proinflammatory mononuclear cells) as well as abnormal neuroplasticity (evidenced by lower dendrite and dendritic spine density and a higher proportion of immature dendritc spine) in aged sickle cell mice compared to age matched controls. We also noted that these neuroimmune and neuronal abnormalities track with the presence and severity of cognitive and neurobehavioral function and that treatment with minocycline (a drug which has been shown to reverse these changes^19, 20, 42, 57^), restored cognitive and neurobehavioral function as well as led to a decrease in evidence of neuroinflammation and abnormal neuroplasticity.

In studies among individuals with SCD, evidence of cognitive and neurobehavioral (such as anxiety) deficits are observed in early childhood^58^ and adolesents.^3, 4, 6^ This, is in addition to the fact that children and adolescents with SCD also experience a high level of silent celerbral infarction (shown to result cognitive deficits)^59^ compared to their non-SCD counterparts,^60-62^ was instrumental in our decision to conduct this study.

Our studies do not show early differences in congitive and behavioural tests (at 6 months of age) but clearly show the presence of an age related decline in cognitive and neurobehavioral function. The reason(s) for this apparent difference between age at development of cognitive decline in sickle cell mice compared to humans is not yet clear to us. However, we recognize that decline in cognitive function is generally an age-related phenomenon^63, 64^ and it has been reported that decline in cognitive function as well as its associated impact in individuals with SCD worsens with age.^65, 66^ Our study was able to capture this aspect of SCD-related cognitive impairment with the emergence of learning, memory, and neurobehavioral deficits by 13 months of age. Thus it was reasonable to conclude that our prior observations (in sickle cell mice) as well as reports in the literature, that cognitive and neurobehavioral abnormalities in SCD increases with age,^14, 67, 68^ or at least partly worsens with age was supported by our results.

It is well established that individuals with SCD are in a state of chronic inflammation and that systemic inflammation is associated with increased vaso-occlussive crisis and stroke. Studies have also indicated that peripheral inflammation is associated cognitive impairment in SCD^2^ as well as neuroinflammation.^19, 22, 37^ In our study, at baseline evaluation of the hippocampus/dentate gyrus revealed significant evidence of neuroinflammation in aged sickle cell mice. This was marked by the higher density of activated microglia and CD45^+^ bone marrow-derived microglia (mononuclear cells) compared to controls (**Fig. 3b i and ii**). The finding of a higher density of bone marrow derived microglia is of particular interest as it shows that in sickle cell disease, mononuclear cells from the peripheral blood migrate into the brain parenchyma in more numbers compared to controls and that these might be contributors to neuroinflammation. Additionally, studies have linked the presence of bone marrow derived microglia in the amygdala and hypothalamus with increased anxiety ^70, 71^ suggesting that levels of bone marrow derived microglia might play a role in the development of the anxiety-like behavior seen in sickle cell disease. Further, we noted that dendritic spine density on pyramidal neurons was lower in aged sickle cell mice compared to controls (**Fig 3c**). Dendritic spine density is a component of neuroplasticity and has been shown to play a significant role in learning and behavior ^72, 73^. Our findings that both neuroinflammation and lower dendritic spine density are potential contributors to cognitive decline in sickle cell disease is strengthened by studies that indicate that the presence and interaction between the two factors are associated with cognitive decline ^74, 75^. These factors provide insight into the cellular and pathobiological mechanisms that contribute to cognitive and neurobehavioral decline in SCD.

Based on the above findings and reports in the published literature, we evaluated whether treatment with minocycline could influence cognitive and neurobehavioral changes as well as the neuronal and neuroinflammatory abnormalities observed in sickle cell mice. Minocycline is a drug (antibiotic) that is capable of crossing the blood brain barrier and that is beneficial to hippocampal integrity^76, 77^. Minocycline has been implicated in the reversal of neuronal cell death and the promotion of neurite outgrowth^39, 43^, reduction of inflammation^78, 79^ and the restoration of cognitive function^42, 80^.

In our study, we first observed that when the sickle cell mice that developed cognitive and neurobehavioral deficits (see **Figs. 3a i-iv**) after being aged from six months (see **Fig 2**) to 13 months of age were randomized to treatment vs. no treatment with minocycline, the treated mice showed a complete reversal of extablished cognitive and neurobehavioral deficits (**Fig S1a i-iii** and **S1b i-iii**). The sickle cell mice that were randomized to not reacived treamtnent with minocylcline did not show any improvement in their performance on cognitive or neurobehavioral tasks. Immunohistochemical analysis also shows that treamtment with minocycline resulted in a decrease in evidence of neuroinflammation indicated by a lower desity of activated microglia as well as CD45^+^ bone marrow-derived microglia (mononuclear cells) in the treated compared to the non-treated sickle cell mice (**Fig. S1c i-ii**).

We repeated the above minocycline treatment study in a larger cohort of mice (N = 15 per group, **Figs 4a** and **4b**) to both confirm our findings and also allowed us to carry out histological analysis to gain more insight into the potential structural changes leading to better cognitive and neurobehavioral function in the minocycline treated mice. Treatment with minocycline as before led to better cognitive and neurobehavioral function in the treated compared to non-treated sickle cell mice. We also observed that sickle cell mice treated with minocycline had signitficantly lower density of activated microglia and CD45^+^ bone marrow-derived microglia (mononuclear cells) in the hippocampus/dentate gyrus compared to non-treated sickle cell mice (**Fig. 4b i and ii)**. Furthermore, treatment of sickle cell mice with minocycline resulted in a neuroplasticity pattern that is similar to that of the healthy controls and thus better than that of non-treated sickle cell mice (**Fig. 5a**). For instance, treated sickle cell mice had significantly higher dendritic spine density (**Fig. 5ai**), but a significantly fewer proportion of immature dendritic spines (**Fig. 5aii**) on hippocampal and cortical pyramidal neurons compared to non-treated sickle cell mice. Additionally, analysis of dendrite aborization shows that sickle cell mice treated with minocycline had more dendrite abors (branches for every 5µm distance away from the soma compared with non-treated sickle cell mice (**Fig. 5b i-iv**).

The results discussed above indicates that cognitive and neurobehavioral deficits in SCD is associated with neuroiflammatory and neuronal pathological changes. Although these are novel observations in SCD, they have been described in non-SCD related cognitive and neurobehavioral deficits. As stated above, it has been shown that cognitive and neurobehavioral deficits as a result of stress from repeat social defeat, is mediated by neuroinflammation seen as increased density of activated microglia and CD45^+^ bone marrow-derived microglia (mononuclear cells) in the hippocampus/dentate gyrus.^19, 20, 22, 81, 82^ The use of minocycline to modify (improve) cognitive and neurobehavioral performance is also not in itself novel. However, this application to SCD-related cognitive and neurobehavioral deficit is the first time it has been used. As shown by our result, treatment of sickle cell mice with minocycline resulted in improvement in cognitive and neurobehavioral function in the treated compared to non-treated sickle cell mice. The mechanisms of this benefit seems to be a modulation of neuroinflammation and neuroplasticity as described in non-SCD models.^19, 42, 83-86^ The sickle cell mice that were treated with minocycline displayed a lower level of cellular evidence of neuroinflammation (CD45^+^ bone marrow-derived microglia (mononuclear cells) in the hippocampus/dentate gyrus), this is thought to be due to the fact that minocycline decreases brain levels of inflammatory cytokines^19^ by inhibiting matrix metalloproteases (MMPs)^84, 87, 88^ and thus prevent the conversion of inactive forms of cytokines and chemokines to active ones. Another mechanism of action of minocycline by which sickle cell mice treated with the drug might benefit, is via inhibition of sphingomyelinase activity^89, 90^ and thus reducing the negative impact of this enzyme on dendrite arborization, and proliferation and maturation of dendritic spines. Finally, it is has also been documenbted that treatment with minocycline results in improved neurogenesis or at least a halting of the gliogenic effects of inflammatory cytokines.^76, 85, 91, 92^ Indeed, despite having similar density of neuron progenitor cells (NPCs or doublecortin and BrDU positive cells) (**Fig S2a**), in our study, we noted a tendency for NPCs in the hippocampus/dentate gyrus of minocycline treated sickle cell mice, to differentiate to mature neurons as opposed to a tendency for differentiation into astroglia (gliogenesis) in the non-treated sickle cell mice (**Fig S2b**)

In conclusion, we have shown that aged sickle cell mice have cognitive and neurobehavioral deficits akin to those described in children and adults with SCD. These deficits worsened with age and were associated with neuroinflammation and abnormal neuroplasticity. Finally, treatment with a neuroinflammation blocking drug such as minocycline, reversed the observed deficits as well as the associated evidence of neuroinflammation and abnormal neuroplasticity. Thus minocycline and/or more novel drugs with smilar mechanisms of action could represent a new line of treatment or prevention paradigm for cognitive and/or neurobehavioral deficits in individuals with SCD. These novel insights into a debilitating sickle cell complication can now be studied futher; investigating the role and interaction of specific cytokines and peripheral and central inflammatory pathways to dissect the pathogenesis and treatment of cognitive and behavioral deficits in individuals with SCD.

## Supporting information

Online-only supplementary material

Fig S1a i-iii

S1b i-iii

## Author Contribution

HIH designed the experiment, HIH, RA, NAR and JJ performed experiments and data analysis, HIH and RA wrote the manuscript and DRA, NAR and JJ provided critical review. HIH and DRA performed final critical review. All authors endorsed the submission of this manuscript.

## Acknowledgement

This study was partially funded by grants to HIH from Emory University Pediatrics Pilot Grant (HeRO) and NIH/NHLBI (U01HL117721, R01HL138423). Also, by the Rodent Behavioral Core (RBC), which is subsidized by the Emory University School of Medicine and is one of the Emory Integrated Core Facilities. Additional support was provided by the Emory Neuroscience NINDS Core Facilities (P30NS055077). Further support was provided by the Georgia Clinical & Translational Science Alliance of the National Institutes of Health under Award Number UL1TR002378. The content is solely the responsibility of the authors and does not necessarily reflect the official views of the National Institutes of Health.

## Data and Laboratory Requests

Request for data used in manuscript can be made to our lab and we will oblige and share our raw data within the limits permissible that will not put us in a disadvantaged position as a laboratory. Mouse models are available from The Jackson Laboratories and we have provided within the manuscript, the sources of our reagents.

