## Supplementary material for "Age and neuroinflammation are important components of the mechanism of cognitive and neurobehavioral deficits in sickle cell disease": Online-only supplementary material

Supplementary Data

**Supplementary Figure 1a.** Sickle (SS) and Control (AA) mice from cohort II that were aged from 6 months (Figure 2) to 13 months were split into three distinct groups. This experiment was to enable us to confirm that memory deficits were age related and to introduce minocycline as a treatment to reduce cognitive and behavioral deficits. These groups consisted of AA (n=19) SS mice (n= 28). A subset of the SS mice was treated with minocycline, creating an SS-treated (n=15) group, while the remaining 13 SS mice were not treated. Novel Object Recognition (NOR) test was conducted to measure behavior and cognition. SS mice exhibited evidence of significant anxiety-like behaviors and depression indicated by the shorter distance traveled (*p<0.0001,* Supplementary Fig 1ai) and thigmotaxis (p=*0.0082;* Supplementary Fig 1aii) when compared to controls. SS-treated mice (72.21% ± 8.37) had an increased preference for novel object when compared to non-treated SS mice (14.86% ± 4.26; *C,* Supplementary Fig 1aiii). SS-treated mice had similar levels of percent preference for novel objects as controls (69.46% ± 4.26).

**Supplementary Figure 1b.** Fear Conditioning was conducted with SS, AA and SS-treated mice tested to assess behavior and cognition. SS mice had no difficulty with fear training but had deficits in learning as indicated by decreased freezing in cued and contextual fear tests. SS, AA and SS-treated mice had no significant difference in the fear training paradigm (Supplementary Fig 1bi) indicated by percent freezing over time. SS (8.47%) and AA (21.8%) mice had similar percent freezing in the contextual fear paradigm but SS-treated (24.83%) mice had higher levels of percent freezing when compared to SS at an early testing time (*p<0.05;* Supplementary Figure 1bii*)*. SS mice had significantly lower percent freezing when compared to AA and SS-treated as time increased. SS mice exhibited a significant decrease in percent freezing in comparison to AA and SS-treated in cued fear context (Supplementary Figure 1bii).

**Supplementary Figure 2a.** The level of proliferating (assessed via BRDU^+^ immunohistochemistry staining) and neural progenitor cells (assessed via DCX^+^ immunohistochemistry staining) in the dentate gyrus were measured in the dentate gyrus of SS, AA, and SS-treated mice. There was no significant difference in proliferating cells per 1000μm^2^ between groups but there was a trend towards SS mice having fewer proliferating cells (10.38± 5.483) when compared to SS-treated (22.58± 12.22) and AA (18.53± 13.80). No significant differences in number of neural progenitor (DCX^+^ cells between genotype or treatment groups; however, similar levels were present in SS (8.86± 4.27) and SS-Treated (8.53± 7.87) mice although, controls or AA mice have significantly more DCX^+^ (20.53 ± 15.96 cells/1000µm) or neural progenitor cells (NPCs). Double labeling with BRDU and DCX was done to determine the number of dividing cells that were neural progenitor cells. While we did not observe any statistically significant differences between groups in the number of cells, there was a trend where AA mice had more BRDU^+^/DCX^+^ or dividing NPCs (21.27 ± 19.49) compared to SS (5.560 ± 2.544) and SS-treated (5.93 ± 5.93).

**Supplementary Figure 2a.** Phenotyping of neural progenitor cells (i.e. DCX^+^ cells) was done with double labeling for young or mature neurons (expressing NeuN^+^) vs. astrocytes (expressing GFAP^+^). There was no statistically significant difference between groups for phenotypically different neural progenitor cells. However, there was a trend towards SS-treated mice having higher numbers (22.70 ± 21.41) of neural progenitor cells that are young or mature neurons (expressing DCX^+^/NeuN^+^) when compared to SS (5.54 ± 4.90) and AA (8.57 ± 6.80) mice. There existed a trend towards SS-treated mice having lower numbers (0.860 ± 0.565) of neural progenitor cells that were astrocytes (expressing DCX^+^GFAP^+^) when compared to SS (3.540 ± 3.540) and AA (8.467 ± 4.619). This results suggests a shift in NPC fate towards differentiation to neurons in sickle cell mice that were compared as opposed to more astrocytes in the sickle cell mice that were not treated.
