## Supplementary figures and images for "Age and neuroinflammation are important components of the mechanism of cognitive and neurobehavioral deficits in sickle cell disease"

### Fig S1a i-iii

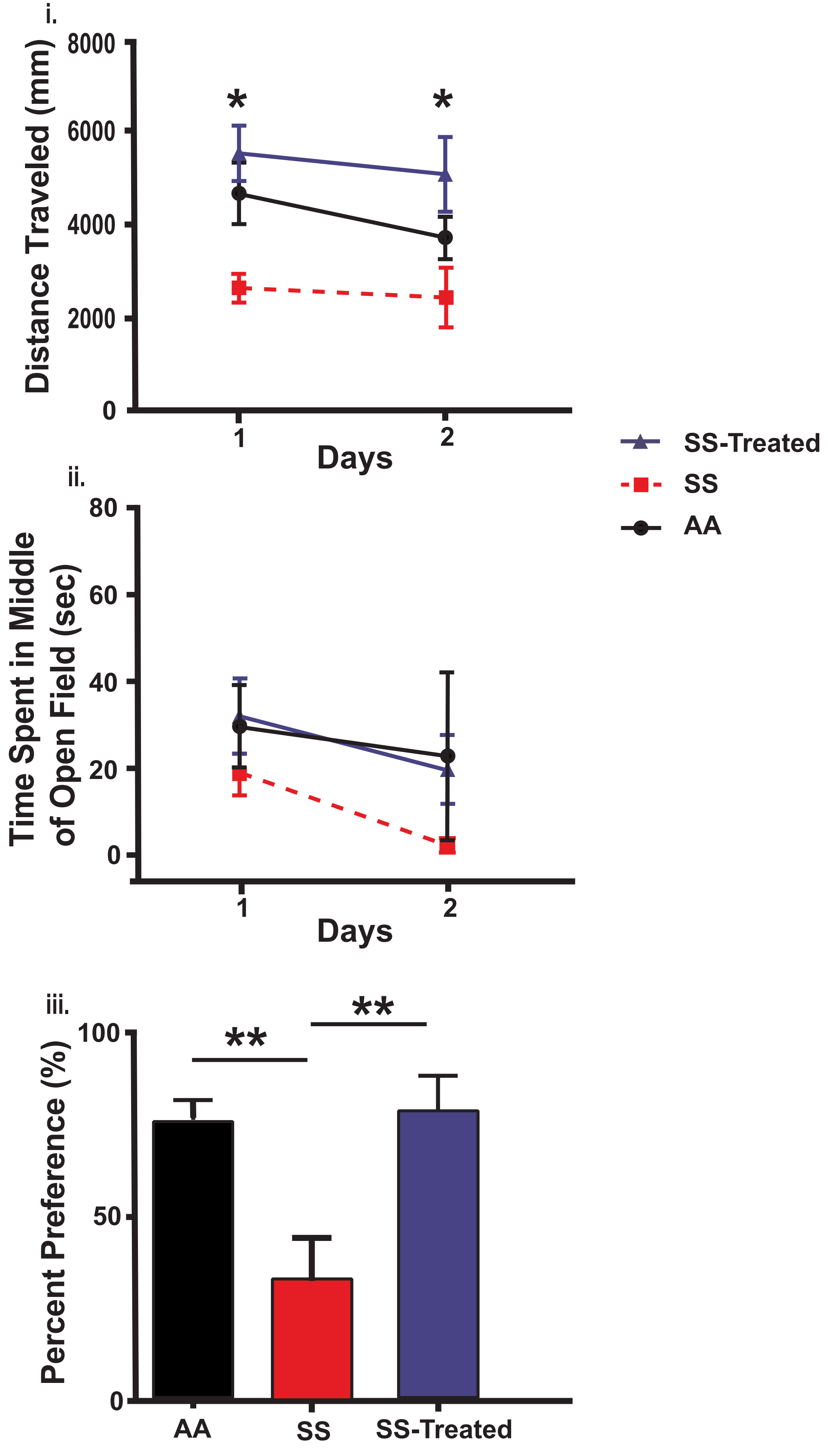


Fig. S1a
Hardy et al

### Fig S2a

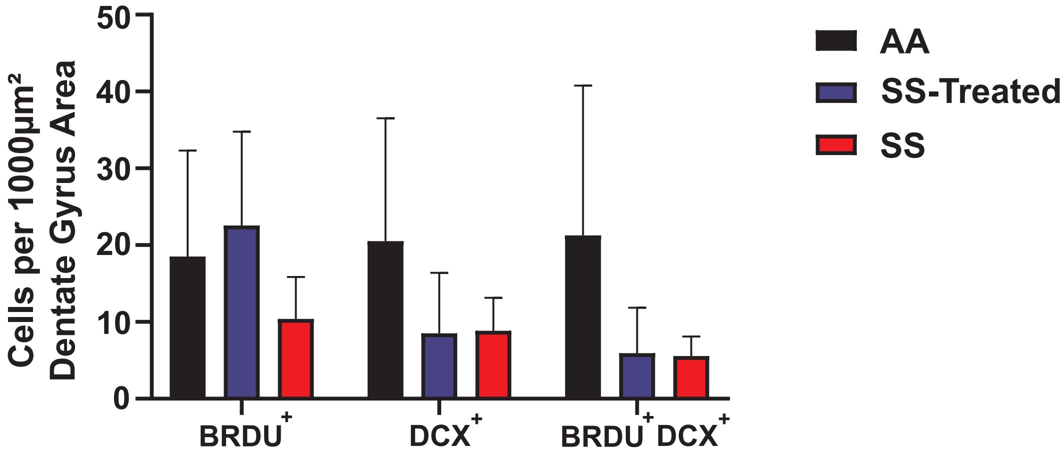


**Fig. S2a**
**Hardy et al**

### Fig S2b

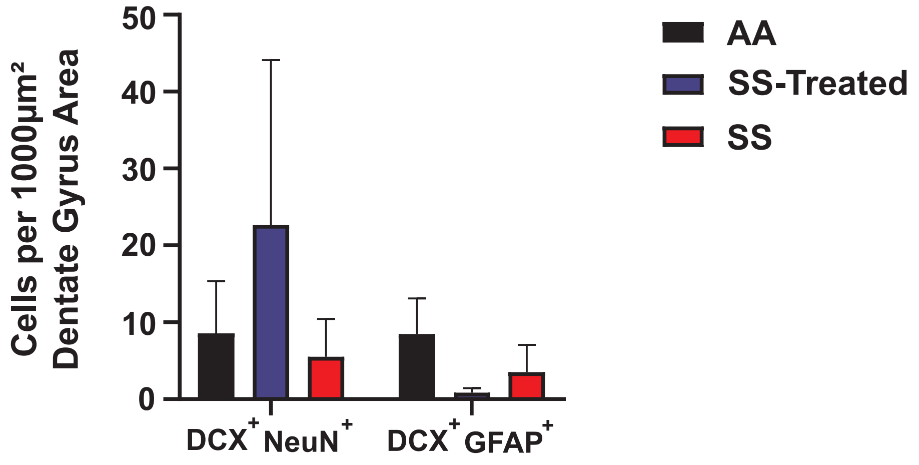


**Fig. S2b**
**Hardy et al.**

### S1b i-iii

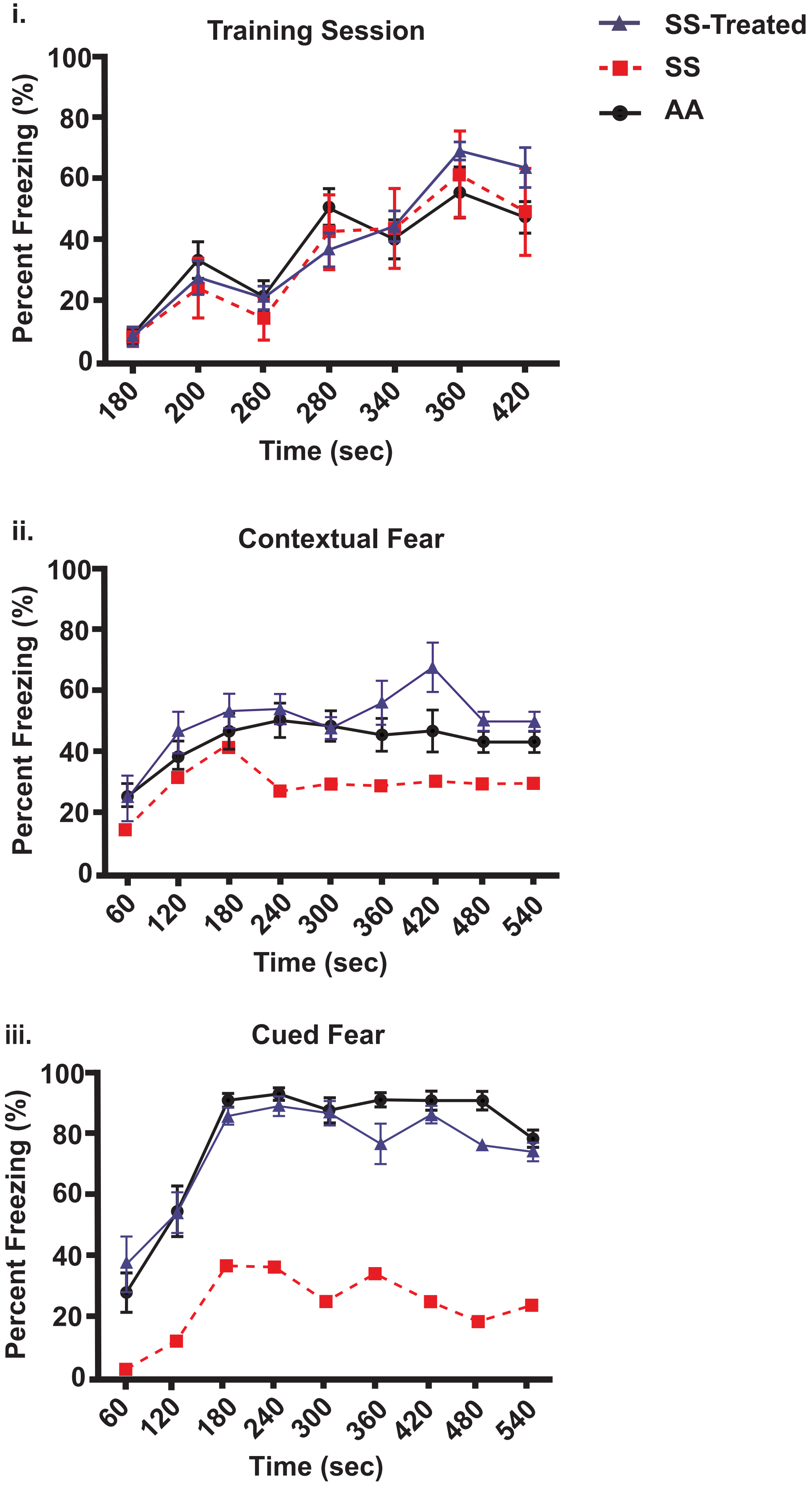


**Fig. S1b**
**Hardy et al**

### S1c i-ii

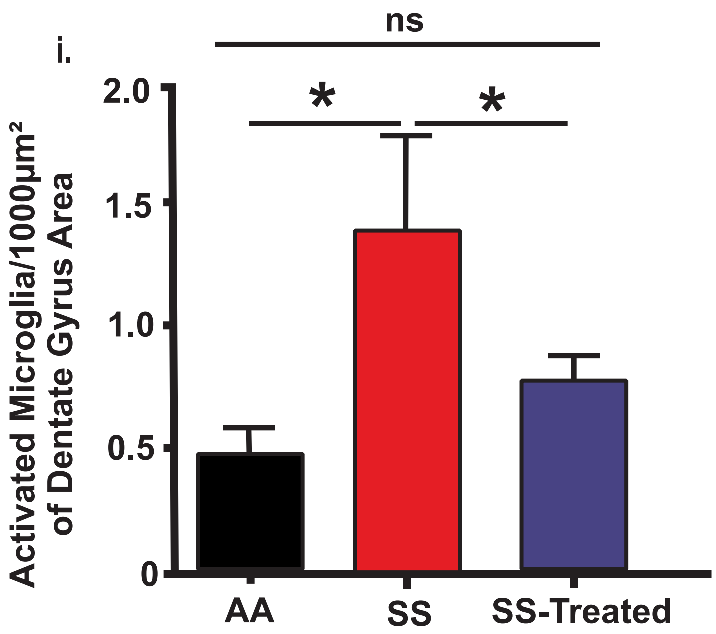


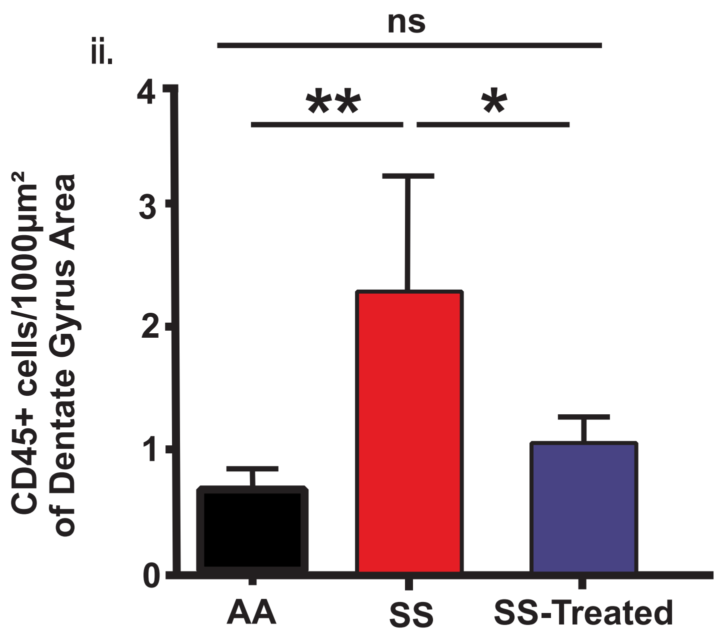


**Fig. S1c**
**Hardy et al**
